# Composition Wheels: Visualizing dissolved organic matter using common composition metrics across a variety of Canadian Ecozones

**DOI:** 10.1101/2020.12.04.412692

**Authors:** Pieter J. K. Aukes, Sherry L. Schiff

## Abstract

Dissolved organic matter (DOM) is a ubiquitous component of aquatic systems, impacting aquatic health and drinking water quality. These impacts depend on the mixture of organic molecules that comprise DOM. Changing climates are altering both the amount and character of DOM being transported from the terrestrial system into adjacent surface waters, yet DOM composition is not monitored as often as overall concentration. Many DOM characterization methods exist, confounding comparison of DOM composition across different studies. The objective of this research is to determine which parameters in a suite of relatively simple and common DOM characterization techniques explain the most variability in DOM composition from surface and groundwater sites. Further, we create a simple visualization tool to easily compare compositional differences in DOM. A large number of water samples (n=250) was analyzed from six Canadian ecozones for DOM concentration, ultraviolet-visible light absorbance, molecular size, and elemental ratios. Principal component analyses was used to identify quasi-independent DOM compositional parameters that explained the highest variability in the dataset: spectral slope, specific-UV absorbance at 255nm, humic substances fraction, and dissolved organic carbon to dissolved organic nitrogen ratio. A ‘Composition Wheel’ was created by plotting these four parameters as a polygon. Our results find similarities in DOM composition irrespective of site differences in vegetation and climate. Further, two main end-member Composition Wheel shapes were revealed that correspond to DOM in organic-rich groundwaters and DOM influenced by photodegradation. The Composition Wheel approach uses easily visualized differences in polygon shape to quantify how DOM evolves by natural processes along the aquatic continuum and to track sources and degradation of DOM.

## Introduction

Dissolved organic matter (DOM) is a pervasive component of aquatic environments and an important determinant of overall water quality and ecosystem function. For example, DOM dictates light, thermal, and pH regimes within lakes [1], complexes with and mobilizes metals [2], and acts as an important redox constituent for biogeochemical reactions [3]. Further, DOM affects drinking water quality through taste, odour, and colour [4], consumption of added oxidizing chemicals, and reactions with chlorine during drinking water treatment processes to form carcinogenic disinfection by-products [5]. The overall reactivity of DOM is determined by its mixture of thousands of organic molecules with differing structural and chemical characteristics. Increased DOM concentrations have been observed in surface waters across the United Kingdom, Europe, and North America [1,6] and linked with declines to water transparency [1,7], that may, in turn have significant effects upon future drinking water treatment options [8,9]. Yet, little information is found on the processes that dictate DOM composition and its impact on the surrounding environment [10]. Hence, quantifying changes to both the amount and composition of DOM across temporal and spatial scales allows for a better understanding of future changes to DOM and its influence upon water quality.

Both concentration and composition of DOM are used to identify the source, quality, and fate of DOM within the environment. The overall DOM concentration is operationally quantified by the concentration of carbon in molecules passing through a chosen filter size (generally between 0.2 and 0.7 μm), whereas DOM composition refers to the mixture of organic molecules that comprise it. Differences in DOM composition have been used to quantify hydrologic mixing and changes to redox potential [11]. Various options are available to characterize DOM composition, but many are expensive, analyze only a subfraction of the DOM, require complicated analysis, or are not widely accessible. For instance, Fourier transform ion cyclotron resonance mass spectrometry (FT-ICR-MS) analyses provides insightful and novel information on DOM composition [12,13] but is not readily available. Alternatively, a number of common and simple measures can provide compositional information on bulk DOM. Ultraviolet and visible light absorbance are used to estimate aromaticity and molecular size of light-absorbing DOM [14–16], while fluorescence provides information on the amount of humic and proteinaceous DOM [17,18]. Simple elemental ratios, such as dissolved organic carbon to dissolved organic nitrogen, indicate DOM lability [19], whereas comparison of various molecular weight groupings can identify differences in DOM sources or processes (i.e. large humic substances from soil-derived sources versus low-molecular weight components from enhanced degradation) [20]. Thus, various characterization techniques can provide a holistic representation of the components that comprise DOM.

Dissolved organic matter is the net product of varying sources and degrees of processing at a point within the watershed. Quantifying changes to DOM composition along a hydrologic continuum can provide information on dominant sources or processes that influence DOM evolution [21]. For instance, DOM sampled from organic-rich substrates typically has high concentrations of large molecular weight, aromatic DOM, while enhanced processing in groundwaters shifts DOM towards smaller components [22–24]. Soil-derived DOM characteristics dominate in headwater streams across various climatic regimes, as shown by higher UV-absorbing properties and higher molecular weight components [18,21]. Conversely, exposure to sunlight and *in-situ* DOM production within surface waters produces DOM with lower UV-absorbing, low-molecular weight components [15]. Once in the aquatic system, differences in DOM composition, or intrinsic controls, are thought to dictate DOM fate more-so than extrinsic controls such as temperature, nutrients, and sunlight exposure [25]. Soil-derived aliphatic components of DOM are preferentially lost as water moves from subsurface to surface waters, at which point aromatic, high-nominal oxidation state DOM becomes further degraded along the fluvial network to more aliphatic, low nominal oxidation state DOM [12,25]. Increased degradation and processing of DOM along the hydrologic continuum reduces its chemodiversity, with persistence of specific components linked to the original DOM composition [25–27]. Hence, compositional measures can be used to better understand how DOM composition varies, and the potential sources or processes that have resulted in the specific mixture of DOM at that point of sampling.

Recent progress in the understanding of DOM and carbon cycling has led to recognition of the importance of characterizing DOM composition along with concentration. Although characterization techniques have improved [28], the final synthesis and discussion of various compositional DOM measures has generally relied on a matrix of scatter plots, requiring the user to jump between graphs and build the comparison themselves. Given the direct impact of DOM composition on water treatment and aquatic health, there is a need to relay compositional information in an efficient and understandable manner to inform policy decisions. This can be difficult if numerous graphs are needed in order to compare DOM differences, all with units and scales specific to the method, especially if the target audience has little knowledge of carbon chemistry or the characterization techniques.

Clear communication of science and its relevance to society is increasing in importance for both informing the general public and supporting evidence-based policy decisions. Effective data visualization can reduce the cognitive load required to understand and retain new knowledge [29,30], creating an accessibility to science that enhances scientific literacy [31–33]. Recently, scientists and graphic designers have collaborated to utilize large datasets and translate data-driven conclusions using novel, creative techniques [34,35]. In terms of DOM composition, graphics have not moved beyond various scatter plots. Further, comparison of different DOM composition metrics have generally focussed on single hydrological environments (i.e. solely lakes [26] or rivers [36]) or on one or two measures across spatial scales or varying environments. Given the complex mixture of organic compounds, and that different measures of DOM composition record different attributes, the combined use of several complimentary measures may provide a better characterization of DOM [37] and enhance public outreach through presentation as a simple, effective image.

The objectives of this study are to select the best suite of readily accessible DOM characterization techniques, and to create a simple, effective visual tool that can be used to illustrate and discuss differences in DOM composition across environmental gradients. These objectives will be accomplished in three parts: 1) determine which broadly-used measures of DOM composition capture the most variability within a dataset of surface and groundwater sites in various Canadian ecozones (groupings of similar areas of biodiversity [38]), 2) create a visualization tool to allow facile comparison of DOM from different samples based upon a set of quasi-independent composition measures, and 3) use the tool to explore DOM composition resulting from differences in hydrological setting and climate across a range of northern environments.

## Methods

### Sites & sampling

Locations were selected where DOM was expected to differ due to differences in surrounding watershed characteristics (e.g. land use, wetland coverage, climate, and vegetation). Various surface and groundwater samples were collected between June to October from the Northwest Territories (Yellowknife, Wekweètì, Daring Lake), high arctic Nunavut (Lake Hazen Watershed), and Ontario (IISD-Experimental Lakes Area) (Fig 1; S1 Table). Surface water samples were collected 0.25 m below the surface from lakes, rivers, creeks, and ponds. Groundwater samples from northern locations (Yellowknife, Wekweètì, Daring Lake, Lake Hazen) were collected from the deepest extent of the mid-summer active-layer, just above the permafrost boundary (0.1 to 0.5 m below surface). Additional samples were collected from Turkey Lakes Watershed (ON), Nottawasaga River Watershed (ON), Grand River (ON), Mackenzie River (NT), and Black Brook Watershed (NB) to expand the ranges of individual composition measures but only overall DOM concentration and select composition metrics are available at these sites (Fig 1; S1 Table). A total of 250 samples were collected and analysed in this study.

**Fig 1.**
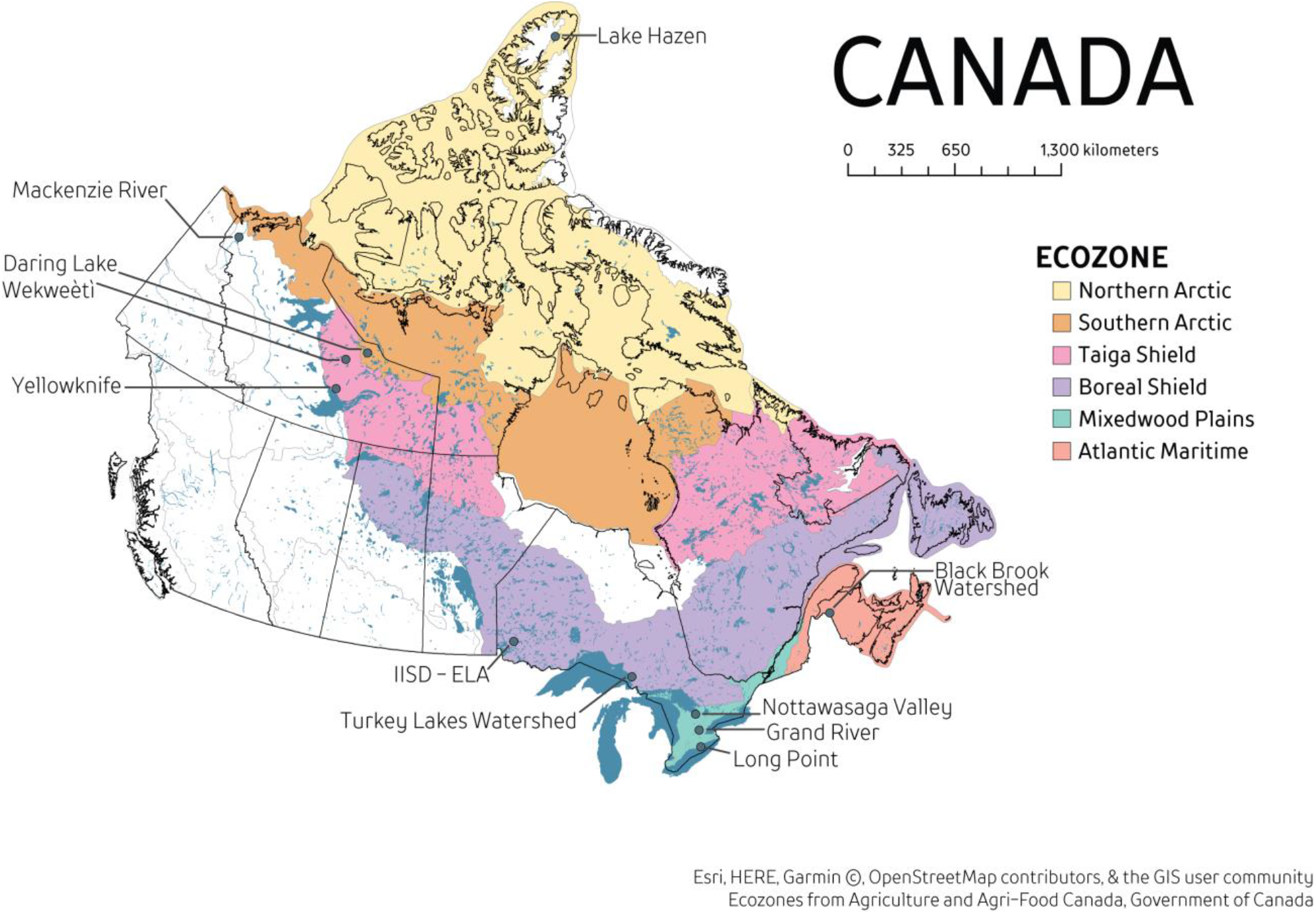
Locations of sampling sites and ecozones. River locations (Grand River, ON; Mackenzie River, NT) are labelled at the mouth of the river.

Surface water samples were collected using a 60 mL syringe and filtered in-field using 0.45 μm syringe-tip filters (Whatman GD/X 45mm) into 40 mL acid-washed, pre-rinsed glass vials. Groundwater samples were collected using a peristaltic pump with an attached 0.45 μm syringe-tip filter. Vials and filters were pre-rinsed with filtered sample water before collection. Samples were kept cool (<4°C) and in the dark until analysis at the University of Waterloo within three weeks of collection.

### DOM quantity & composition analyses

Dissolved organic carbon and total nitrogen concentrations were measured using a Shimadzu Total Organic Carbon (TOC-L) Combustion Analyzer with TNM-1 module. Dissolved organic nitrogen (DON) was calculated as the difference between total dissolved nitrogen concentration and the sum of inorganic nitrogen species (nitrate, nitrite, and ammonium). Inorganic nitrogen species were measured using SmartChem 200 Automated Chemistry Analyzer (Unity Scientific, MA United States). The DOC:DON ratio was calculated using molar concentrations of DOC (M_C_) and DON (M_N_).

Absorbance was measured in a 1 cm cuvette using a Cary 100 UV-VIS Spectrophotometer (Agilent, CA United States) at 5 nm increments from 200 to 800 nm. Deionized water was used to zero the instrument and run intermittently during analyses to correct for baseline drift. The Naperian absorption coefficient (*a*; m^-1^) was calculated using:

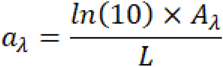

where *A* is the baseline-corrected absorbance at wavelength λ and *L* is the cell length (m). A suite of absorbance characteristics were then calculated (Table 1).

**Table 1.**
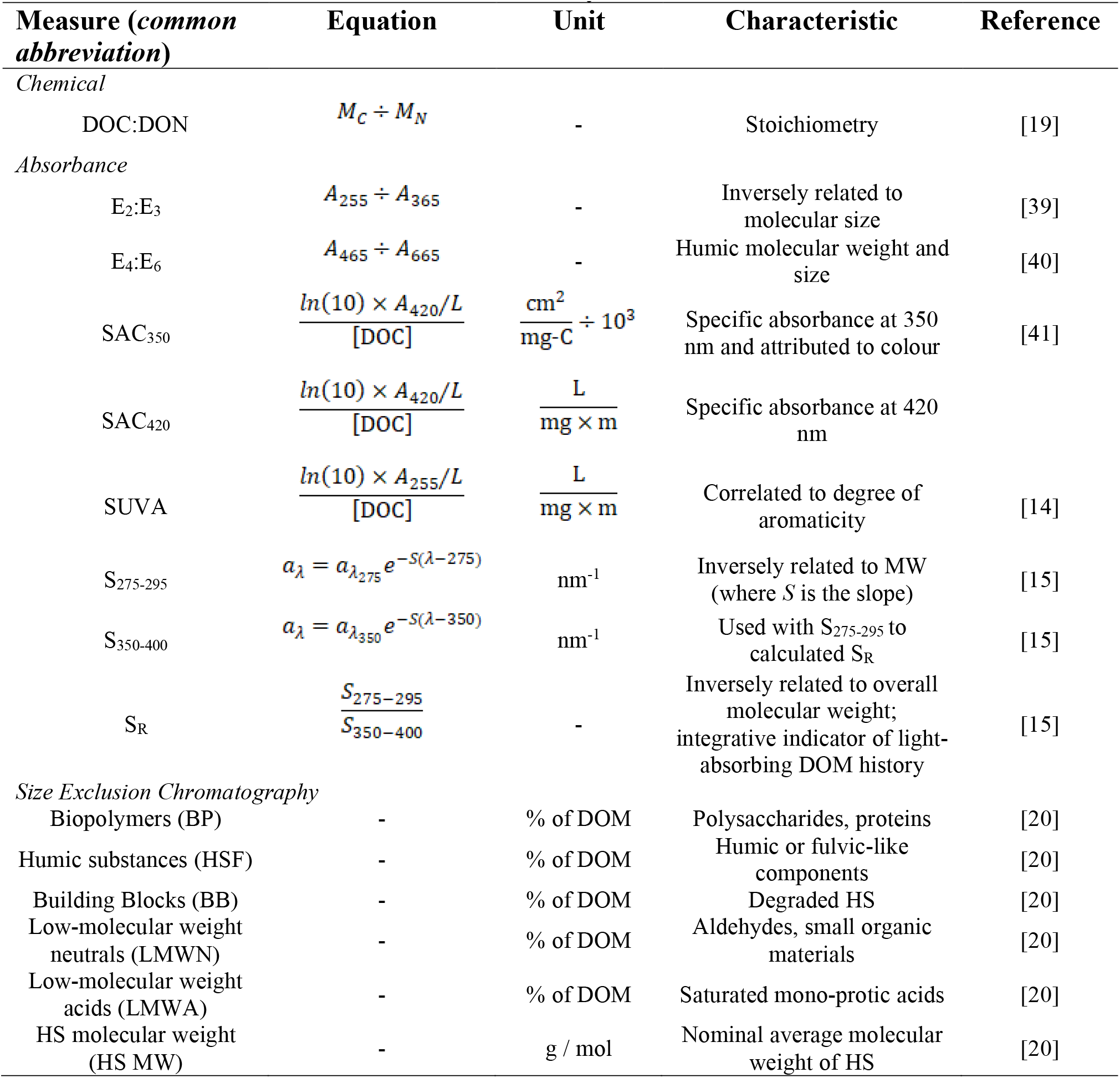
**Dissolved organic matter composition as described by chemical, absorbance, and molecular-size based measures used in this study**.

Molecular-size based fractions of DOM were determined using a size exclusion chromatography technique (Liquid Chromatography – Organic Carbon Detection, LC-OCD) at the University of Waterloo. Detailed instrument setup and analysis is described elsewhere [20]. Briefly, the sample was injected through a size-exclusion column (SEC; Toyopearl HW-50S, Tosoh Bioscience) that separated DOM based on hydrodynamic radii into five hydrophilic fractions (from largest to smallest): biopolymers (BP; polysaccharides or proteins), humic substances fraction (HSF; humic and fulvic acid fraction), building blocks (BB; lower weight humic substances), low molecular weight neutrals (LMWN; aldehydes, small organic materials), and LMW-acids (LMWA; saturated mono-protic acids). A portion of the sample by-passes the SEC for determination of the overall DOC concentration, here referred to as DOM concentration in mg C/L. A number average molecular weight was derived only for the HSF based on elution time. Duplicates run at six concentrations yield a precision for the LC-OCD of <0.1 mg C/L for all fractions. Concentrations of each fraction were calculated using specialized software (ChromCALC, DOC-Labor, Germany) that integrated chromatograms from the LC-OCD.

### Statistical analyses & Composition Wheel design

Samples from sites with multiple sampling events were averaged to create a single value per site. Data were analysed using unconstrained ordination analysis via principal components analysis (PCA) on a subset of samples that contained all composition measures listed in Table 1 (subset n=130). Data were scaled before PCA and analysed using R Statistical Software [42].

The Composition Wheel (CW) is a polygon drawn from axes of various composition measures that are independent of DOM concentration in order to focus solely on differences in DOM composition. Composition Wheel parameters were chosen based on the highest contribution of variables explaining the first two principal component axes (S1 Fig). Further, independent measures of DOM composition were preferentially chosen to minimize overlap in information between similar techniques. Each CW axis corresponds to a specific parameter. For each axis, the individual value for each sample is normalized as a value between the maximum and minimum encountered for that parameter within the dataset. All R code and data used to create the DOM CW, manuscript figures, and statistical analyses can be found at https://github.com/paukes/DOM-Comp-Wheel.

## Results

### DOM concentration & composition

DOM concentrations ranged from 0.1 to 273 mg C/L, with highest mean values in boreal groundwater, pond, and creek samples (Fig 2; S1 File). The highest DOM concentrations were found in groundwater environments in Yellowknife, while the lowest concentrations were found in high arctic environments (Fig 2). High arctic seeps, rivers, and lakes also generally had the lowest average DOC:DON values, but higher average specific ultra-violet absorbance at 255 nm (SUVA), spectral slope (S_275-295_), and HSF values than other locations. SUVA and spectral slope values covered the known range as described broadly in the literature (SUVA: 1.1 to 21 L/(mg·m); spectral slope: 0.005 to 0.032 nm^-1^). Highest SUVA values (>11 L/(mg·m)) were found at two sites: the anoxic bottom of a boreal lake and in the high arctic groundwater, but absorbance may be affected by high iron concentrations [14,43]. These two sites have unusually high iron concentrations and were not included further in the discussion. Values of DOC:DON ranged between 9 to 124, and were lowest in rivers and high arctic samples. Humic substances fraction (HSF) ranged from 14% to 85% and on average were lowest in lakes. Overall, groundwater samples contained the highest DOC:DON and HSF values, and lowest S_275-295_. Hence, both DOM concentration and composition varied across geographic scales and hydrological environments.

**Fig 2.**
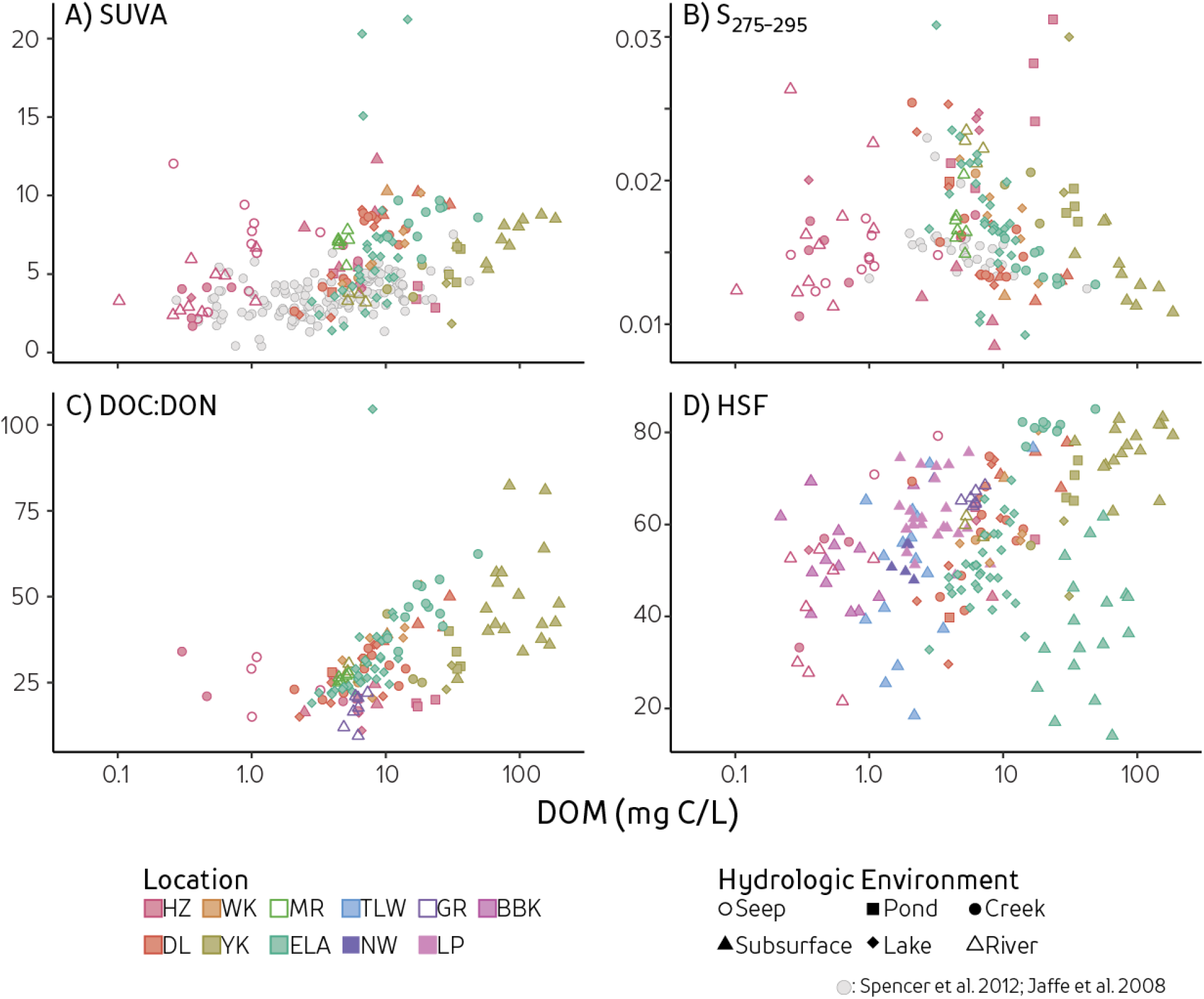
Compositional measures versus total DOM concentration. Measures include A) SUVA, specific ultraviolet absorbance at 255 nm, B) S_275-295_, spectral slope between 275 to 295nm, C) DOC:DON, and D) HSF; humic substances fraction. Colours represent geographical sampling locations (HZ: Lake Hazen Watershed, NU; DL: Daring Lake, NT; WK: Wekweètì, NT; YK: Yellowknife, NT; MR: Mackenzie River, NT; ELA: IISD-Experimental Lakes Area, ON; TLW: Turkey Lakes Watershed, ON; NW: Nottawasaga River Watershed, ON; GR: Grand River, ON; LP: Long Point, ON; BBK: Black Brook Watershed, ON) while shapes represent hydrologic environments. Light grey circles represent two other published DOM characterization studies conducted at similar scales [16,44].

### PCA on DOM composition measures

The first three principal component axes explained 66% of the variance in DOM composition using the measures contained in the dataset, with PC1 and PC2 accounting for 54% of the variability (Fig 3). Comparison of the first two principal components (PC) yield four distinct groupings of strongly-contributing measures: I) SUVA, SAC_420_, SAC_350_; II) HSF, HS MW; III) S_275-295_, E_2_:E_3_, and S_R_; and IV) BB, LMWN, and BP. Groups I and II were negatively associated to groups III and IV. Highest contributions to PC1 and PC2 axes were HSF, SAC_350_, SAC_420_, SUVA, and S_275-295_. Variables with contribution to PC1 and PC2 lower than 2% were E_4_:E_6_, DOC:DON, S_350-400_, and LMWA (Fig. S1). Absorbance parameters normalized to DOM concentrations (SAC_350_, SAC_420_, and SUVA) all plotted closely to each other and trended positively with measures of HSF, HSF molecular weight, and DOC:DON. Absorbance techniques plotted perpendicular to LC-OCD size fractions (Fig 3).

**Fig 3.**
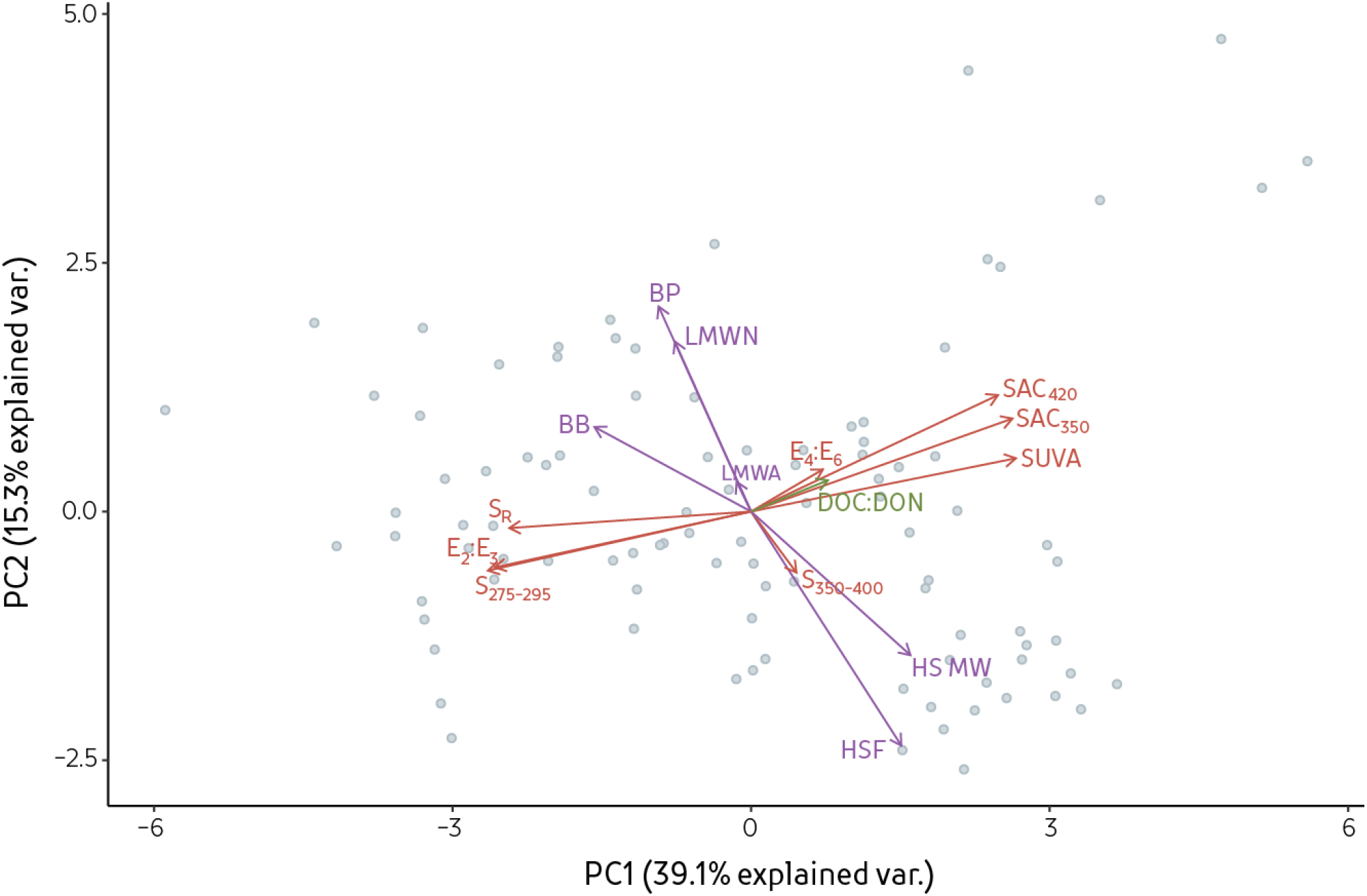
Principal component analyses for samples from different ecozones. Grey dots represent individual samples, while vectors represent absorbance (red), elemental (green), and LC-OCD (purple) compositional measures.

Based on contribution to the first two PC axes, we selected four DOM composition measures to define DOM composition. Further, these selected measures are quasi-independent as they are based on different analytical principles or attributes (SUVA, S_275-295_: two different absorbance aspects; HSF: size-exclusion chromatography; DOC:DON: stoichiometry). Although the absorbance at different wavelengths within a sample are closely related [21], they can be used to provide different information on DOM components as one characterizes the amount of UV-absorbing components normalized to DOM concentration at a specific wavelength (SUVA) while the other is associated with the shape of the absorbance spectra and varies with the distribution of DOM molecular weights (S_275-295_). These four measures include three high-contributing variables to PC1 and PC2 axes (HSF, SUVA, and S_275-295_). The length of the CW axis, and thus CW size, is dictated by the range of each compositional measure encountered in the entire dataset. Composition Wheels are then used as a basis for comparing DOM.

### Shapes of Composition Wheels

Different environments are characterized by CW different shapes. Ponds and lakes tend to have lower values of SUVA and DOC:DON and higher S_275-295_, resulting in a triangle shape elongated to the top-right corner (Fig 4). Groundwater DOM samples form a trapezoidal shape due to higher HSF and SUVA values than other sites. Differences in shape are found within locations between different hydrologic settings whereas similarities in shape are found within hydrological setting across locations. For instance, Yellowknife pond sites are similar in shape to groundwater DOM, while the creek, lake, and river shapes reflect DOM with higher S_275-295_, lower DOC:DON, and lower HSF. Representation of different measures using these shapes allows for a facile comparison of DOM composition among samples.

**Fig 4.**
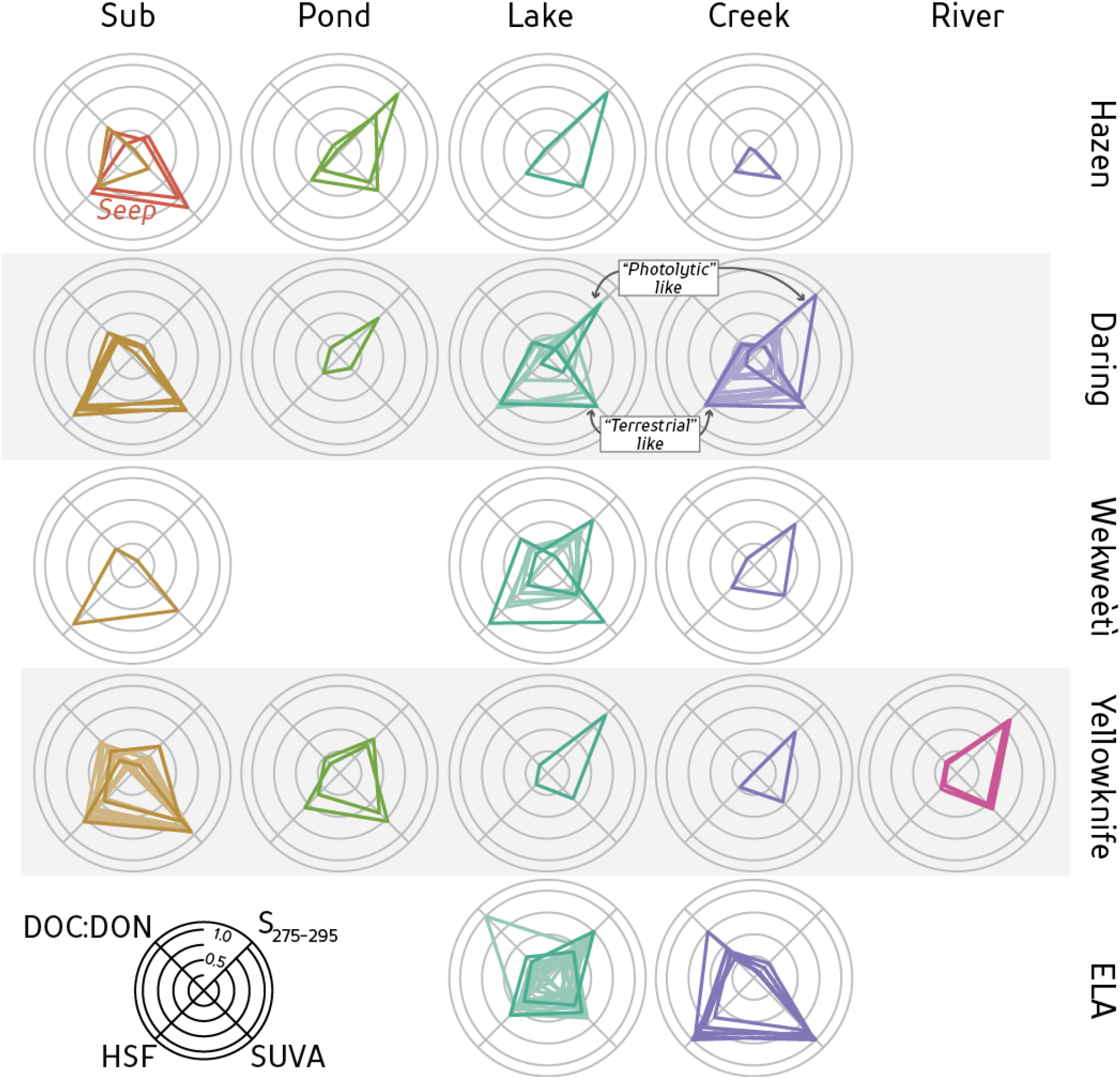
Composition wheels for DOM from different hydrologic settings within each geographical site. Axes are a numerical value normalized relative to the maximum and minimum encountered within the dataset for each parameter. Parameters for each axis found in bottom left. The orientation of the composition wheel (CW) axes are arbitrary. Different samples from the same hydrological and geographic setting are plotted within the same CW. Two ‘end member’ compositions are highlighted in sites with many samples to help visualize the continuum of DOM mixtures: groundwater-like DOM (low DOC:DON, high SUVA and HSF) or photolyzed DOM (high S_275-295_).

Shapes also help to visualize and highlight the most extremes in DOM composition for each location. The spread of compositions is not evenly distributed between such extreme shapes at any given site but sampling was not targeted *a priori* based on shape. Most surface water sites contained a range in DOM composition between these extremes, forming either shapes similar to groundwater DOM or thin, elongated triangles in the S_275-295_ direction. For this reason, extremes were highlighted in sites with many different shapes to help identify a continuum of possible DOM sources or mixtures (Fig 4). Further, some sites did not conform to these end-members, as seen by the ELA creek sample with high DOC:DON, allowing easy identification of anomalous data or sites that may warrant further investigation.

## Discussion

### Comparison of DOM composition measures

Ranges in individual measures of DOM composition are large at both low and high concentrations of DOM (Fig 4, S2 Table) making it difficult to consistently separate sites or hydrological environments across different measures. For instance, accounting for both DOM concentration and DOC:DON may help differentiate some locations, but the range in DOC:DON values across moderate DOM concentrations covers most sites (Fig 2). Hence, new techniques that simultaneously consider different aspects of DOM composition, and are independent of concentration, are needed to compare variations in DOM composition across sample types and environments.

Different characterization techniques provide different information on DOM. In particular, the perpendicular relationship between absorbing and non-absorbing parameters observed among the first two PC axes indicate a range in different DOM properties are captured. For instance, all ELA creeks contain high and similar HSF values, yet we can use absorbance techniques based on molecular weight or UV-absorbing capability (via SUVA; Fig 2) to differentiate between these samples. However, correlations between size-based and absorbance parameters do not mean causation. For example, higher SUVA values are not consistently related to HSF proportions even though this is often assumed based on the higher aromaticity and UV-absorbing capabilities of humic material [14,45]. Daring Lake creeks have high SUVA but low HSF. Although absorbance parameters are easily measured and helpful in comparing different DOM compositions, they do not accurately reflect non-absorbing components (e.g. stoichiometry or functional groups) that could differ among samples.

### Effects of DOM Processes and CW End Members

Similar DOM CW shapes were found within hydrologic settings even though samples were collected from sites with varying climates and vegetation, spanning areas of boreal shield watersheds (IISD-ELA) to high arctic (Lake Hazen). Further, the range in values across the four CW measures compared well with DOM from other studies (i.e. similar DOM concentrations and SUVA values to rivers in the United States [16,44] and Canadian boreal lakes [46]) indicating that DOM composition and CW shape may not be unique to its locale. The similarity in CW shapes across different locations may result from analogous drivers of DOM fate. For instance, the proportion of biodegradable DOM was found to be a function of DOM composition and nutrients, rather than ecosystem or region-specific characteristics [47], suggesting that differences in the quality of DOM may not result from location alone. Hence, CWs can be used to amalgamate different compositional measures and quantify similarities in DOM sources and processing across a range of geomorphic, climate, and vegetation environments.

Composition Wheels can be further used to determine the degree of DOM degradation and facilitate comparisons of change along the aquatic continuum. This requires knowledge of the effect of processing, as well as the end-members (e.g. original sources of DOM). Microbial degradation of DOM induces a shift towards higher SUVA and lower DOC:DON (Fig 5; [48–51] but not S_275-295_, resulting in a distinctive effect of microbial DOM transformation on the CW. Photolytic processing of DOM results in an increase in S_275-295_ values and a general decrease in the other three DOM quality measures [52–54]. Further, photolytic degradation affects all four compositional axes, whereas microbial degradation only affects two axes (Fig 5), indicating that certain measures of DOM composition respond differently to different DOM degradation processes. This has important implications for studies using single-characterization techniques as some parameters may not always faithfully serve as useful surrogates for others. For instance, microbial degradation changes SUVA but not HSF (Fig 5), even though SUVA has been linked to DOM aromaticity in humic substances [14]. Thus, comparative differences in a CW can help determine the dominant processes that produce the observed DOM composition of a sample. Further, knowing the degree DOM composition changes with photolysis allows CW to be used in a quantitative manner to quantify the amount of degradation. Although the quantitative response of DOM sources to photo- or microbial degradation is not yet sufficiently known across different environments, in areas where these rates of change to individual measures have been measured, CWs can be used in a quantitative manner to assess the extent and major processes contributing to DOM loss.

**Fig 5.**
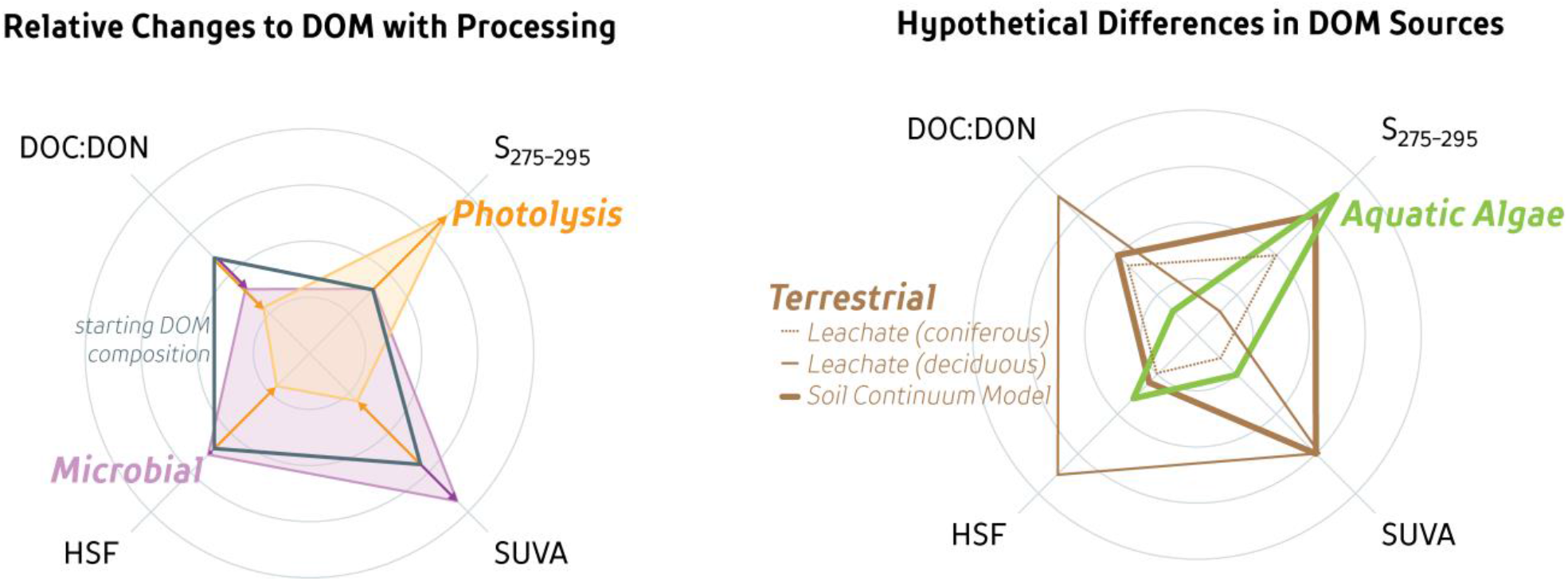
Changes to a Composition Wheel resulting from photolytic and microbial degradation (left) and differences in terrestrial versus aquatic DOM sources (right). Changes to dissolved organic matter (DOM) (grey) with photolytic (orange; 12 to 18 day exposure to sunlight) or microbial (purple; 30-day experiment) degradation based on experimental incubations of natural DOM samples [51] (left). The mean decrease in DOC concentration was 18% during photolysis and 10% during microbial degradation experiments (n=11, 9; respectively). Hypothetical differences in DOM sources (right) are based on various leachate experiments in literature [22,37,55–59].

Laboratory experiments are useful to assess DOM source composition prior to extensive biotic or abiotic processing. Although this study did not conduct analysis on leachates, various studies have used similar DOM composition measures to identify DOM leachates. Leachates (organic matter steeped in de-ionized water) result in either low molecular weight, low aromatic DOM from mosses and coniferous sources [22,55] or large, humic, aromatic DOM components from other plant material and deciduous leaf litter [37,57,58]. DOM produced from aquatic (primary production) consists of low molecular weight, high protein and nitrogen content, and low UV-absorbing components and is thought to be rapidly consumed [37,56,59,60]. These parameters can also be represented by a CW (Figure 5b – using a range in values based on what is presented in literature), and provide a shape to compare DOM origins from aquatic primary production versus terrestrial plants.

Two dominant DOM compositional end-member shapes are evident in our dataset: groundwater DOM sources and DOM altered by photolysis. Other shapes are intermediate between these two distinctive end-members. Degradation in groundwater in organic-rich substrates allows for the accumulation of large, aromatic components of DOM (high HSF, SUVA, and low S_275-295_; [61]), resulting in a large trapezoid shape CW. This trapezoid shape is consistent for groundwater DOM across most sites, supporting previous groundwater studies that attribute the narrow DOM compositional range to groundwater processing [23]. In contrast, a ‘kite-like’ shape results from the effects of photolysis (Fig 4), resulting in DOM with lower molecular weight and less UV-absorbing components [15]. Further, photolysis end-member CWs are similar in shape to the end-products of photolytic experiments (Fig 5; [51]), and are clearly seen in surface waters across all sites (Fig 4). Hence, these end-member shapes may constrain the possible range in CW shapes that can be found within aquatic environments.

The groundwater/photolysis end-member categorization frames our conceptual model (Fig 6) that quantifies how the two end-member CW shapes evolve depending on the aquatic-terrestrial hydrologic connection (Fig 6; pathway A-b) or exposure to sunlight (Fig 6: pathway A-B). For instance, creeks with little to no processing of groundwater-derived DOM can be identified by CW shapes similar to groundwater DOM, as seen in ELA and some Daring creeks (Fig 4). ELA lakes have a wide gradient of CW shape and thus overall DOM quality (Fig 4). The use of CW to pair process-based knowledge of DOM composition with hydrologic transport provides a framework that can be used identify key knowledge gaps (i.e. rates of degradation, kinetics based on composition, relative organic source contributions, water residence time) and quantify the relationship between DOM sources and degree of processing within the environment.

**Fig 6.**
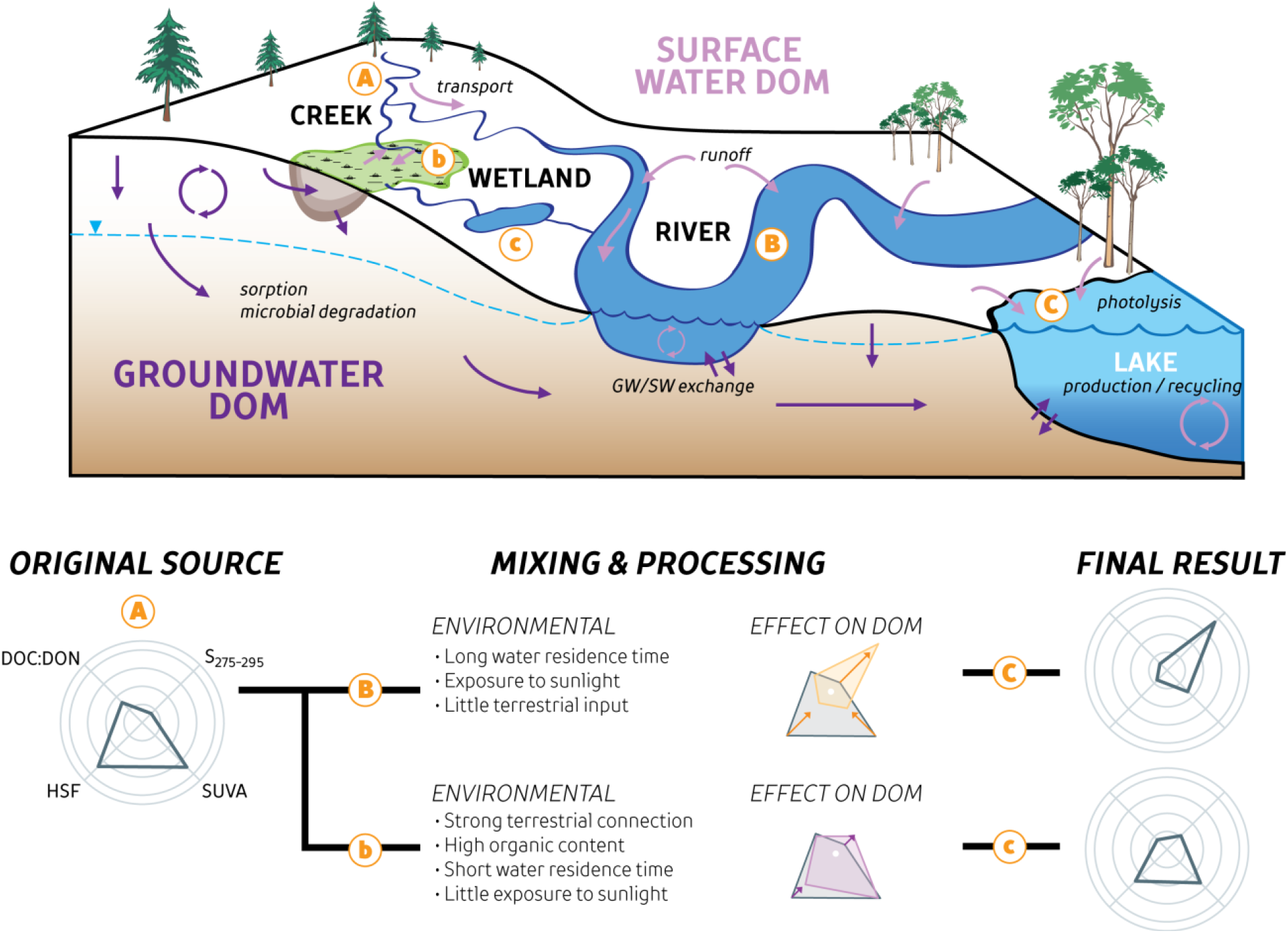
Conceptual model of DOM evolution along the aquatic continuum using Composition Wheels. A conceptual model of a few select processes and sources of dissolved organic matter (DOM) along the aquatic continuum. Two main hydrologic flowpaths are illustrated (A-B-C and A-b-c) and originate from the same source ‘A’. However, each flowpath undergoes different processing within the environment. Arrows represent the transport or addition of DOM among groundwaters (dark purple) and surface waters (light purple), while lowercase text highlights some of the specific processes and sources encountered. The wetland ‘b’ represents an area of organic matter degradation and DOM accumulation within organic-rich groundwaters, resulting in the transport of large, aromatic DOM components into surface water ‘c’. Conversely, DOM is rapidly transported down the river ‘B’ and into a large lake ‘C’, resulting in much higher exposure to sunlight. The use of Composition Wheels to illustrate differences in sources and processing is shown on the bottom panel, comparing DOM from an organic-rich source with longer exposure to sunlight (A-B-C) versus the same DOM with less sunlight exposure and greater microbial degradation (A-b-c).

Certain creeks and rivers from YK or WK do not fit into the end-member categories as they contain shapes with larger right sides (higher S_275-295_ and SUVA than other samples), indicating other processes may be important in determining DOM composition. This may be a result of our sampling of mostly oligotrophic systems that are dominated by terrestrial carbon. We hypothesize that these different shapes represent surface water systems with greater *in-situ*, or autochthonous, contributions of DOM, generally characterized by higher amounts of proteins and smaller molecular weight components [62–64]. Thus, although our dataset was not comprehensive in terms of capturing the full range of DOM sources, the CW-informed conceptual model provides an initial framework to build the factors leading to a specific DOM composition.

### Adaptability of DOM Composition Wheels

Expressing DOM composition with only four concentration-independent parameters excludes advantages of other techniques not used in this study. However, agreement between multiple parameters allows for selection of surrogates for the CW. LC-OCD is not as widely available but many studies have traditionally used resins [16,65,66] or other size-exclusion columns [67,68] to characterize DOM. Further, LC-OCD fractions of humics have been well correlated to measures of ^13^C-NMR and fluorescence measures such as HIX [12,69,70]. Although fluorescence was not used in this study, strong associations between certain fluorescence parameters and absorbance or molecular-weight groupings could be used to replace LC-OCD defined fractions, such as using PARAFAC modelling to discern components most similar to HSF [12,70] (Fig 7). These fluorescence parameters are also independent measures of DOM composition and could be readily substituted into the CW. Elemental ratios of DOC:DON are generally positively correlated with humic-like fluorescence and negatively correlated to protein-like components [71]. Other absorbance indices such as slope ratio and E_2_:E_3_ (Table 1) are potential surrogates for S_275-295_ (Fig 2), providing various parameters that can be substituted for one another when comparing studies where only one of these measures is provided. Measurements of SUVA could be substituted with SAC_420_, SAC_350_, or E_4_:E_6_ [25,71]. However, although SUVA has been correlated with HIX and to fluorescence component C3 [25,71], the opposite has also been found [70]. Composition Wheels can easily incorporate advanced techniques, such as FT-ICR-MS, that can provide information on the molecular heterogeneity of DOM, identification of either terrestrial-like (via condensed and polycyclic aromatics or polyphenolic compounds) or *in situ* DOM sources (higher aliphatic and peptide-like compounds), or to identify differences in H:C and O:C due to microbial or photodegradation [12,13,72]. Associations between different characterization techniques allow for mixing of techniques and comparison of different variables, indicating a wide-range of applicability of CW within environmental sciences.

**Fig 7.**
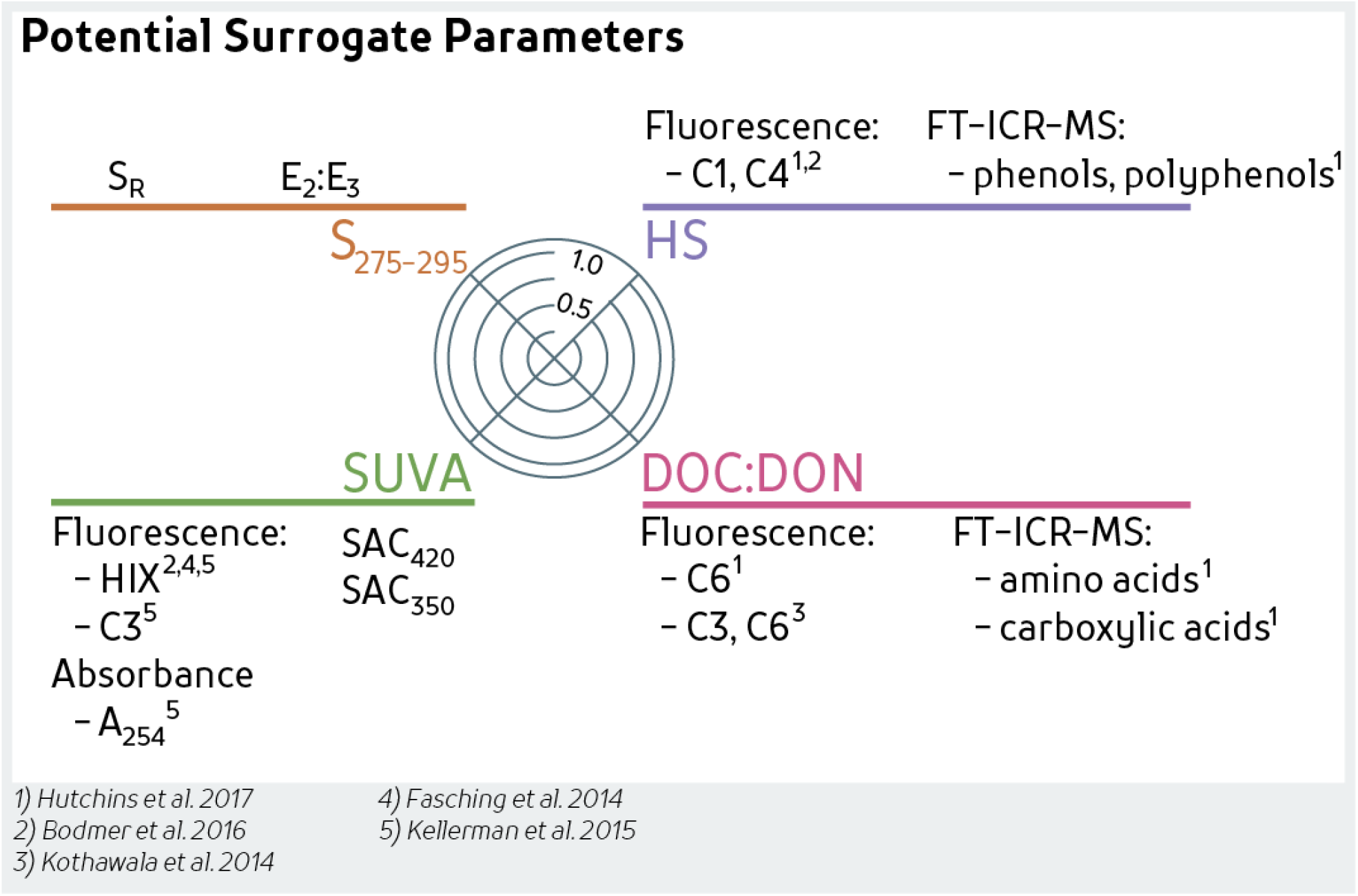
Surrogate parameters for the DOM Composition Wheel.

The CW visualization method provides an efficient communication tool not only among scientists, but also between scientists and other stakeholders concerned with water quality, including community members. Shapes from CWs show clearly how DOM differs, whereas changes in the numerical value of multiple metrics are only easily grasped by those with familiarity with the methods and environmental ranges. Further work could include pairing different shapes of DOM with key DOM roles such as disinfection demand and by-product formation, metal mobility, and mercury bioaccumulation. By reducing the complexity of independent DOM measures and creating an easily comparable shape, differences in DOM composition can be easily grasped and communicated to larger audiences.

## Acknowledgements

We thank Michael English, Roy Judas, and Jordan Reid for field assistance in collecting samples in the Northwest Territories; Paul Dainard, Kyra St. Pierre, Igor Lehnherr, Vince St. Louis, Jessica Serbu, Maria Cavacao, Victoria Wisniewski, and Michael English for field assistance at Lake Hazen, NU; Will Robertson for assistance at Long Point, ON; Environmental Geochemistry Lab, University of Waterloo for sampling the Grand River, ON; John Spoelstra for sampling Black Brook Watershed, Nottawasaga River Watershed, and Turkey Lakes Watershed. We also thank the Community of Wekweètì and the Wek’èezhìi Land & Water Board for their support, and the Community-Based Monitoring Program and Department of Environment and Natural Resources, Government of Northwest Territories, for collection of samples from the Mackenzie River, NT. We would like thank Richard J. Elgood for logistical, field, and technical help, Monica Tudorancea assistance with the LC-OCD analysis, and Mackenzie Schultz for help with Figure 1. We are grateful for the high arctic logistic support provided by Parks Canada, and funding from the Polar Continental Shelf Program (PCSP), Northwest Territories Cumulative Impact Monitoring Program, Northern Scientific Training Program, and the National Science and Engineering Research Council of Canada. We thank Igor Lehnherr, Susan Ziegler, Chris Parsons, Roland Hall, and three anonymous reviewers for their comments on earlier drafts of this manuscript. PJKA also benefitted from scholarships from the Garfield Weston Foundation, TD Friends of the Environment, and the University of Waterloo. Data used in this manuscript can be found in the publicly-available Wilfrid Laurier University Scholars Portal Dataverse (https://doi.org/10.5683/SP2/NG6D02), while all R code used for statistical analyses, figure creation, and a Composition Wheel template is available from https://github.com/paukes/DOM-Comp-Wheel.

## Supporting information

**S1 Fig.**
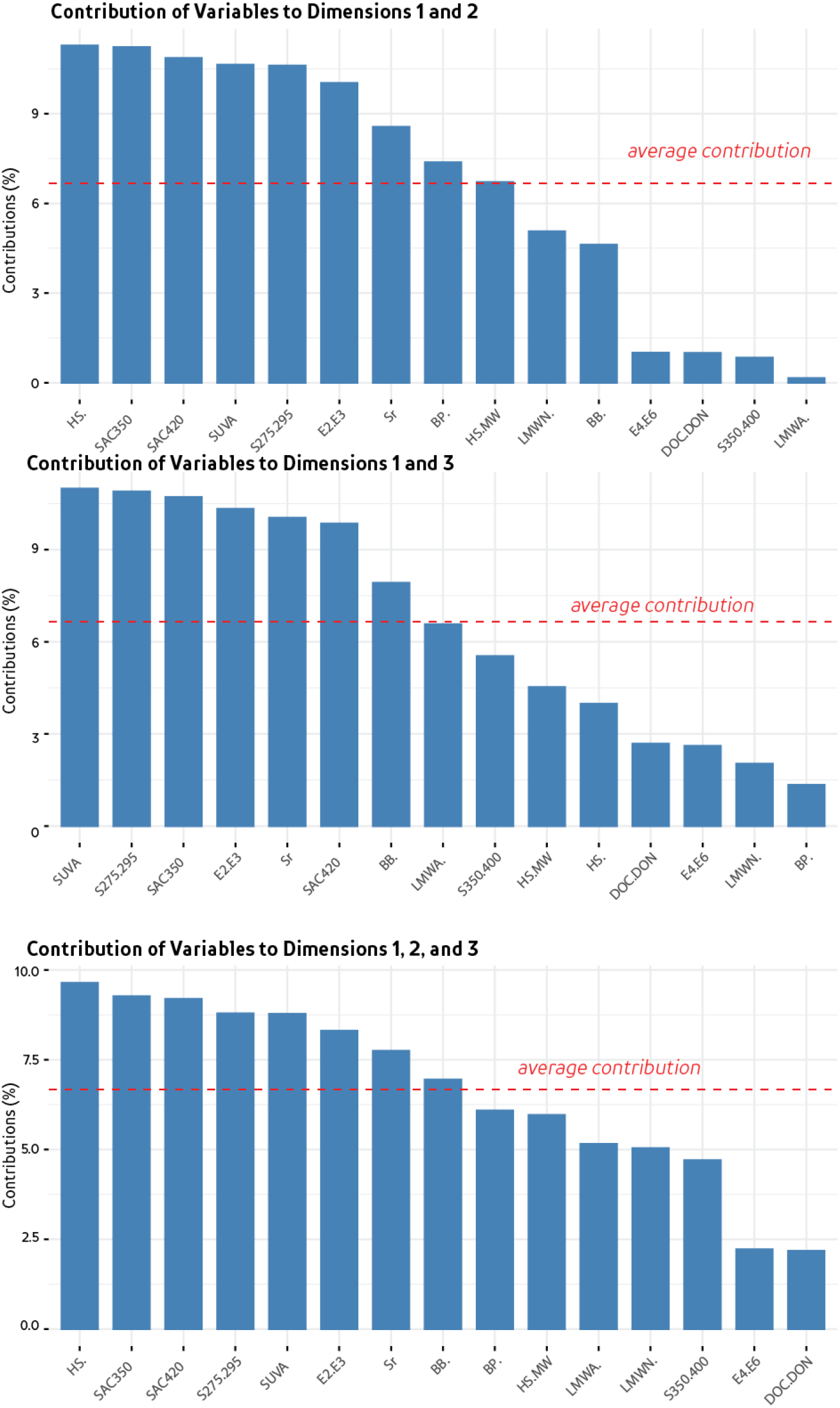
Contribution of each variable within the PCA.

**S1 Table.**
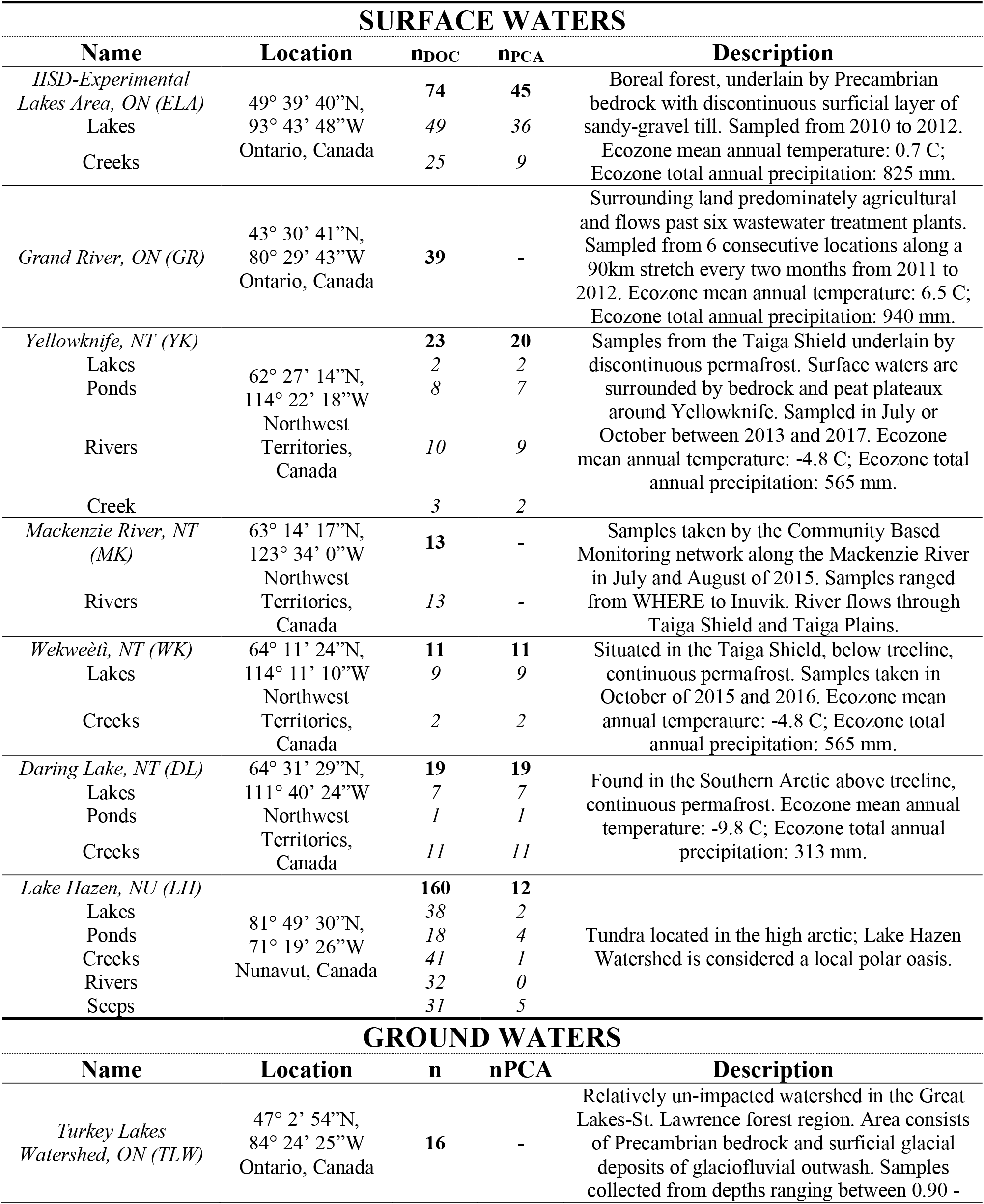
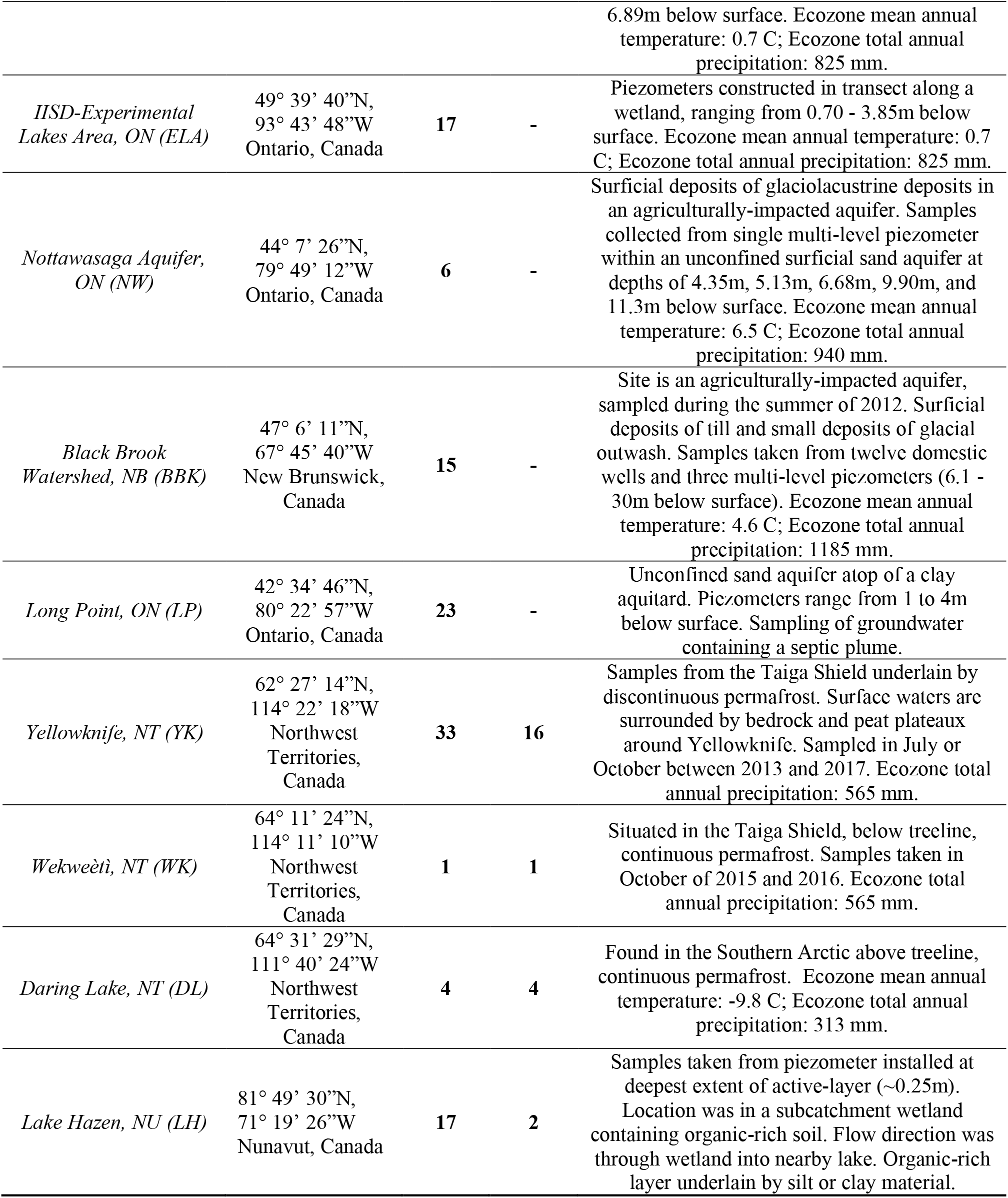
Environmental description for all sampling sites. Included are the number of total samples taken for DOM concentration with at least one compositional measure (n_DOC_), and number of samples used from that site in the PCA analysis (n_PCA_).

**S2 Table.** DOM Composition versus Concentration Statistics. Results from a linear model of dissolved organic matter (DOM) composition (SUVA, slope between 275-295nm, DOC:DON, and humic substances fraction) as predicted by the overall DOM concentration (mg C/L; log transformed).

| Parameter | Linear Regression to log(DOM) |  |
| --- | --- | --- |
|  | R<sup>2</sup> | p-value |
| SUVA | 0.10 | <0.01 |
| S275 | 0.01 | 0.61 |
| DOC:DON | 0.33 | <0.01 |
| HSF | 0.09 | <0.01 |

